# Lymphopenia-induced T cell proliferation is a hallmark of severe COVID-19

**DOI:** 10.1101/2020.08.04.236521

**Authors:** Sarah Adamo, Stéphane Chevrier, Carlo Cervia, Yves Zurbuchen, Miro E. Raeber, Liliane Yang, Sujana Sivapatham, Andrea Jacobs, Esther Bächli, Alain Rudiger, Melina Stüssi-Helbling, Lars C. Huber, Dominik J. Schaer, Bernd Bodenmiller, Onur Boyman, Jakob Nilsson

## Abstract

Coronavirus disease 2019 (COVID-19), caused by infection with severe acute respiratory syndrome coronavirus 2 (SARS-CoV-2), has a broad clinical presentation ranging from asymptomatic infection to fatal disease. Different features associated with the immune response to SARS-CoV-2, such as hyperinflammation and reduction of peripheral CD8^+^ T cell counts are strongly associated with severe disease. Here, we confirm the reduction in peripheral CD8^+^ T cells both in relative and absolute terms and identify T cell apoptosis and migration into inflamed tissues as possible mechanisms driving peripheral T cell lymphopenia. Furthermore, we find evidence of elevated serum interleukin-7, thus indicating systemic T cell paucity and signs of increased T cell proliferation in patients with severe lymphopenia. Following T cell lymphopenia in our pseudo-longitudinal time course, we observed expansion and recovery of poly-specific antiviral T cells, thus arguing for lymphopenia-induced T cell proliferation. In summary, this study suggests that extensive T cell loss and subsequent T cell proliferation are characteristic of severe COVID-19.

## Introduction

The global epidemic of severe acute respiratory syndrome coronavirus 2 (SARS-CoV-2), which is the causative agent of coronavirus disease 2019 (COVID-19), has affected at least 15 million people worldwide and led to over 600,000 deaths as of July 2020. Patients with COVID-19 have a wide spectrum of symptoms ranging from asymptomatic infection to severe acute respiratory distress syndrome (ARDS) (Arons *et al.*, 2020)(Tong *et al.*, 2020)(Li *et al.*, 2020)(Wu and McGoogan, 2020)(Yang *et al.*, 2020). Advanced age and comorbidities are risk factors for the development of severe disease (Zhou *et al.*, 2020). Furthermore, individuals with severe disease have increased amounts and longer duration of SARS-CoV-2 viral shedding in the respiratory mucosa and of viral RNA in blood as compared to individuals with mild COVID-19 (Liu *et al.*, 2020)(Zheng *et al.*, 2020)(Hadjadj *et al.*, 2020). Several reports have demonstrated associations between severe disease and elevation of systemic inflammatory markers such as C-reactive protein (CRP), procalcitonin, and IL-6 (Feng *et al.*, 2020)(Ruan *et al.*, 2020)(Wu *et al.*, 2020)(Zhou *et al.*, 2020). Taken together these data suggest that inefficient antiviral immunity and ensuing hyper-inflammation might underlie the pathogenesis of severe COVID-19.

T cells are central players in antiviral immunity. Effector T cells eliminate virus-infected cells and assist in innate antiviral responses. Virus-specific T cells also support B cells responses, which culminate in the production of virus-specific antibodies (Cervia *et al.*, 2020). It has been convincingly shown that severe COVID-19 is associated with reduced amounts of CD3^+^ T cells in peripheral blood and that the extent of the T cell decrease correlates with disease severity (Hadjadj *et al.*, 2020) (Feng *et al.*, 2020)(Diao *et al.*, 2020)(Du *et al.*, 2020). The reduction in peripheral T cells appears to be particularly prominent within the CD8^+^ T cell compartment, but it remains unclear if this is due to trafficking of CD8^+^ T cells into tissues with ongoing SARS-CoV-2 replication, increased elimination of CD8^+^ T cells during COVID-19, or pre-existing abnormally low levels of CD8^+^ T cells in individuals who experience severe disease.

Recent reports examining tissue-resident lymphocytes, including CD8^+^ T cells, in patients with severe COVID-19 suggest involvement of both migration and apoptosis, but these studies have evaluated only small numbers of patients (Xu *et al.*, 2020)(Liao *et al.*, 2020)(Chen *et al.*, 2020). The reduced amount of CD8^+^ T cells in patients with severe COVID-19 has also been associated with increased expression of surface markers typical of T cell exhaustion, such as PD-1 and TIM-3 (Diao *et al.*, 2020)(Schultheiss *et al.*, 2020). Increased frequencies of exhausted T cells have been reported in patients with viral infections who have persisting viremia and inefficient clearance of virus-infected cells (Mueller and Ahmed, 2009)(Shin and Wherry, 2007) but have also been associated with insufficient type I interferon responses (Kolumam *et al.*, 2005) and subsequent lack of efficient CD4^+^ T cell help in acute infections (Wang *et al.*, 2012).

To facilitate a detailed investigation of the peripheral T cell compartment, we performed mass cytometry, flow cytometry, targeted proteomics, and functional assays to phenotypically and functionally characterize the changes associated with symptomatic COVID-19 and relate them to disease severity. We observed global T cell loss, especially among CD8^+^ T cells, which was associated with disease severity and increased T cell apoptosis, exhaustion, and functional impairment. Furthermore, we observed signs of lymphopenia induced proliferation and expansion of poly-specific T cells, suggesting that T cell loss and the compensatory non-specific proliferation are central to the pathogenesis of severe COVID-19.

## Methods

### Subjects and clinical data

Patients 18 years and older with symptomatic, RT-qPCR confirmed SARS-CoV-2 infection were recruited at four different hospitals in Zurich, Switzerland. Both hospitalized patients and outpatients were recruited into the study and all participants gave written informed consent. The study was approved by the Cantonal Ethics Committee of Zurich (BASEC 2016-01440).

Peripheral blood was obtained from the 70 patients at the time of inclusion into the study and standardized clinical data were collected longitudinally for all included patients. Two patients with severe COVID-19 had hematological malignancy and were excluded from further analysis. Two patients presented with an atypical disease course with complete symptom resolution and a long symptom-free interval, followed by development of respiratory symptoms and PCR positivity so that re-infection could not be excluded, and were also excluded. Disease severity was classified according to the WHO criteria (World Health Organization, 2020) and patients were stratified as having mild disease, including mild illness (n=20) and mild pneumonia (n=10), and severe disease, including severe pneumonia requiring oxygen therapy (n=21) as well as mild (n=7), moderate (n=8), and severe ARDS (n=4). Of the blood samples from the 66 subjects, 98%, 85%, 94%, and 98% were analyzed by routine flow cytometry, mass cytometry, targeted proteomics, and functional T cell assays, respectively. Additionally, 22 healthy donors without evidence of previous symptomatic SARS-CoV-2 were also included as controls. Of the blood samples from the 22 controls, 95%, 100%, 76%, and 90% were analyzed by routine flow cytometry, mass cytometry, proteomics, and functional T cell assays, respectively.

### Blood collection and sample processing

Venous blood samples were collected in BD vacutainer EDTA tubes unless otherwise specified. Whole blood was centrifuged, and plasma was removed and frozen at −80 °C. The remaining blood was diluted with PBS and layered into a SepMate tube (STEMCELL, catalog number 85460) filled with Lymphodex solution (Inno-Train Diagnostik GmBH, catalog number 002041500). After centrifugation, peripheral blood mononuclear cells (PBMCs) were collected and washed with PBS. An aliquot of PBMCs was processed in preparation for mass cytometry. Briefly, 1×10^6^ PBMCs were centrifuged and resuspended in 200 μl of 1.6% paraformaldehyde (Electron Microscopy Sciences) in RPMI 1640 medium and fixed at room temperature for 10 min. The reaction was then stopped by adding 1 ml of 0.5% bovine serum albumin and 0.02% sodium azide in PBS. Cells were centrifuged and stored at −80 °C after disruption of the cell pellet. The remaining PBMCs were resuspended in 10% dimethyl sulfoxide and frozen at −80 °C.

### Flow cytometry

For quantification of the main T cell subsets, blood samples were processed in the accredited routine immunology laboratory at the University Hospital Zurich. Briefly, 10 μl of Krome Orange anti-CD45 (Beckman Coulter, clone J33), APC-A750 anti-CD3 (Beckman Coulter, clone UCHT-1), APC anti-CD4 (Beckman Coulter, clone 13B8.2), and AF700 anti-CD8 (Beckman Coulter, clone B9.11) were added to 100 μl of whole blood in a 5-mL round-bottom tube, and samples were incubated at room temperature for 15 min. After adding 2 mL of VersaLyse solution (Beckman Coulter, catalog number A09777) with 2.5% Fixative Solution (Beckman Coulter, catalog number A07800), samples were further incubated for 15 min at room temperature. Immediately prior to data acquisition, 100 μl aliquots of Flow-Count Fluoroshperes (Beckman Coulter, catalog number 7547053) were added to each sample. Data were acquired on a Navios flow cytometer and analyzed using the Kaluza software.

CD127 levels were quantified in samples from six healthy donors. PBMCs were obtained as described above, and 1×10^6^ cells from each donor were incubated with BUV 737 anti-CD3 (BD Biosciences, clone UCHT1), BUV 395 anti-CD4 (BD Biosciences, clone SK3), AF700 anti-CD8 (Biolegend, clone 5C3), PECy5 anti-CD25 (Invitrogen, clone 61D3), BV510 anti-CD45RA (Biolegend, clone HI100), BV421 anti-CCR7 (Biolegend, clone G043H7), PE-Dazzle anti-CD127 (Biolegend, clone A019D5) at room temperature for 15 min. Cells were then washed with staining buffer (PBS, 2% FBS, 2 mM EDTA) and acquired on a BD Fortessa. Data were analyzed with the Flow-jo software.

### Mass cytometry analysis

#### Mass cytometry barcoding

As references for the CyTOF analysis, PBMCs from a healthy donor were divided into three aliquots and stimulated with 0.1 μg/mL phytohemagglutinin for 24 h, 1 μg/mL lipopolysaccharide plus 1.5 μg/mL monensin for 48 h, or left unstimulated. After treatment, the PBMCs were fixed and frozen as described above.

Homogenous staining was ensured by barcoding of 1×10^6^ cells from each patient following a barcoding scheme consisting of unique combinations of four out of eight barcoding reagents as previously described (Zunder *et al.*, 2015).

Six palladium isotopes (^102^Pd, ^104^Pd, ^105^Pd, ^106^Pd, ^108^Pd, and ^110^Pd, Fluidigm) were chelated to 1-(4-isothiocyanatobenzyl)ethylenediamine-N,N,N’,N’-tetraacetic acid (Dojino), and two indium isotopes (^113^In and ^115^In, Fluidigm) were chelated to 1,4,7,10-tetraazacy-clododecane-1,4,7-tris-acetic acid 10-maleimide ethylacetamide (Dojino) following standard procedures (Zivanovic, Jacobs and Bodenmiller, 2014).

Mass-tag barcoding reagents were titrated to ensure an equivalent staining for each reagent; final concentrations were between 50 nM and 200 nM. Cells were barcoded using the previously described transient partial permeabilization protocol (Behbehani *et al.*, 2014). Samples were randomly loaded into wells of 96-well plates and were analyzed in two independent experiments. Three reference samples were loaded on each plate. Cells were washed with 0.03% saponin in PBS (PBS-S, Sigma Aldrich) and incubated for 30 min with 200 μl of mass tag barcoding reagents diluted in PBS-S. Cells were then washed three times with cell staining medium (CSM, PBS with 0.5% bovine serum albumin and 0.02% sodium azide), and samples from each plate were pooled for cell staining with the antibody panel.

#### Antibodies and antibody labeling

The antibodies used in this study are listed in Table S1. Antibodies were labeled with the indicated metal tags using the MaxPAR antibody conjugation kit (Fluidigm). We assessed the concentration of each antibody after metal conjugation using a Nanodrop (Thermo Scientific) and then supplemented each antibody with Candor Antibody Stabilizer. Inventories of antibodies used in this study were managed using the cloud-based platform AirLab (Catena *et al.*, 2016).

#### Staining with antibodies and data acquisition

Prior to staining, pooled cells were incubated with FcR blocking reagent (Miltenyi Biotec) for 10 min at 4 °C. Cells were stained with 400 μL of the antibody panel per 10^7^ cells for 45 min at 4 °C. Cells were then washed three times in CSM, once in PBS, and resuspended in 0.4 ml of 0.5 μM nucleic acid Ir-labeled intercalator (Fluidigm) and incubated overnight at 4 °C. Cells were then washed once in CSM, once in PBS, and once in water. Prior to acquisition, cells were diluted to 0.5 x 10^6^ cells/mL in Cell Acquisition Solution (Fluidigm) containing 10% of EQ™ Four Element Calibration Beads (Fluidigm). Samples were acquired on an upgraded Helios CyTOF 2. Individual .fcs files collected from each set of samples were pre-processed using a semi-automated R pipeline based on CATALYST to perform individual file concatenation, normalization, compensation, debarcoding, and batch correction (Fig. S1).

#### Data analysis

T cells from each sample were classified using a random forest classifier (R package randomForest), as described in (Chevrier *et al.,* 2020). To visualize the high-dimensional data in two dimensions, the t-SNE algorithm was applied on data from a maximum of 1,000 randomly selected cells from each sample, with a perplexity set to 80, using the implementation of t-SNE available in CATALYST (Nowicka *et al.*, 2017). All antibody channels were used to calculate the t-SNE. Data were displayed using the ggplot2 R package or the plotting functions of CATALYST (Nowicka *et al.*, 2017).

To visualize marker expression on t-SNE maps, data were normalized between 0 and 1 with a maximum intensity defined as the 99th percentile. For hierarchical clustering, pairwise distances between samples were calculated using the Spearman correlation or euclidean distance, as indicated in the figure legend. Dendrograms were generated using Ward.2’s method. Heatmaps were displayed using the pheatmap package. Clustering analysis of the was performed using the R implementation of PhenoGraph run on all samples simultaneously, with the parameter *k*, defining the number of nearest neighbors, set to 100 (Levine *et al.*, 2015).

### Flow cytometric assay for specific cell-mediated immune responses in activated whole blood

Venous blood was collected in sodium heparin tubes. Blood was diluted with RPMI 1640 medium supplemented with 10% FBS, 100 IU/mL penicillin and 100 IU/mL streptomycin (all from Gibco). Blood cells were stimulated with pokeweed mitogen, concanavalin A, *Staphylococcus* enterotoxins A and B, or antigens from varicella zoster virus (VZV), adenovirus, cytomegalovirus (CMV), herpes simplex virus 1 (HSV1), or herpes simplex virus 2 (HSV2) or left unstimulated for 7 days. Cells were then stained with live/dead fixable Aqua stain (Thermo Fisher, catalog number L34957) and with Cyto-stat tetrachrome (FITC anti-CD45, PECy5 anti-CD3, PE anti-CD4, and ECD anti-CD8, Beckman Coulter, catalog number 660713). Unstimulated cells and pokeweed mitogen-stimulated cells were additionally stained with AF750 anti-CD19 (Beckman Coulter, clone J3119). Samples were incubated 15 min at room temperature, then washed once with blocking buffer (PBS, 1% bovine serum albumin, 2 mM EDTA), resuspended in VersaLyse solution (Beckman Coulter, catalog number A09777) with 2.5% Fixative Solution (Beckman Coulter, catalog number A07800) and incubated 15 min at room temperature. After an additional wash with blocking buffer, cells were resuspended in blocking buffer. Data were acquired on a Navios flow cytometer and analyzed with Kaluza analysis software. Net stimulation was calculated by subtracting the percentage of CD3^+^ blasts or CD19^+^ blasts over all lymphocytes in the unstimulated sample from the percentage of CD3^+^ blasts or CD19^+^ blasts over all lymphocytes in stimulated samples.

### Cytokine measurements

Serum was collected in BD vacutainer clot activator tubes (Becton Dickinson). The samples were processed in the accredited immunological laboratory at the University Hospital Zurich. IFN-γ and TNF-α were quantified using R&D Systems ELISA kits.

### Proteomics analyses

Heat-inactivated plasma samples were analyzed using the Olink® Proteomics 92-plex inflammation immunoassay. In brief, binding of paired oligonucleotide-labeled antibodies directed against the targeted serum proteins led to hybridization of the corresponding DNA oligonucleotides, which were extended by a DNA polymerase. Expression was quantified using real-time PCR. The values are reported as log2-scaled normalized protein expression (NPX) values. Only data on samples that passed the quality control are reported. Samples below the detection limit are reported as the lower limit of detection.

### Statistics

Descriptive statistics for the study cohort (stratified by healthy, mild disease, and severe disease) are presented as median and interquartile ranges unless otherwise specified. For comparisons of two independent groups the Wilcoxon-Mann-Whitney test was used. The p values were adjusted for multiple comparisons with the Holm method. For comparisons of more than two independent groups the non-parametric Kruskal-Wallis test was used. The Fisher’s exact test was used for categorical variables. A linear regression model was used to quantify the relationship between variables. Significance was assessed by non-parametric methods unless otherwise specified. All tests were performed two sided. Analyses were performed with R (version 4.0.0) and with Graph Pad Prism.

## Results

### Characteristics of COVID-19 patients and healthy subjects

To characterize the immune response associated with SARS-CoV-2 infection we conducted a prospective observational study on symptomatic COVID-19 patients recruited from hospitals in the Canton of Zurich, Switzerland. The included patients were stratified based on clinical disease severity at the time of sampling (mild, n=28; severe, n=38); a group of healthy controls was included for comparison (n=22) (**Fig. 1a**). Patients were sampled at a single time point during their symptomatic phase, with approximately 70% sampled within 18 days of symptom onset (**Fig. 1a**). Standard laboratory parameters and clinical characteristics of the included patients are presented in **Table 1**. In agreement with previous studies (Mathew *et al.*, 2020)(Schultheiss *et al.*, 2020), we observed that patient age was positively correlated with disease severity (**Fig. 1b**) and that males were overrepresented in the severe COVID-19 subgroup (**Fig. 1c**).

**Table 1.** Clinical and laboratory characteristics of healthy subjects and COVID-19 patients.

|  | Healthy | Mild cases | Severe cases |
| --- | --- | --- | --- |
| Number at sampling | 22 | 28 | 38 |
| Median age (median (IQR<sup>a</sup>) [yrs]) | 32.50 (29.25-48) | 50.5 (34.50-60.25) | 67.5 (59.0-79.0)** |
| Gender (m/f) | 11/11 | 12/16 | 24/14 |
| Time since symptom onset (days) | - | 12.86 ± 10.71 | 20.21 ± 11.96* |
| **COVID-19 disease severity sampling / max severity<sup>b</sup>** |  |  |  |
| Mild illness– no. | - | 18 / 16 | - |
| Mild pneumonia – no. | - | 10 / 8 | - |
| Severe pneumonia – no. | - | - | 20 / 19 |
| Mild ARDS – no. | - | - | 7 / 7 |
| Moderate ARDS – no. | - | - | 7 / 8 |
| Severe ARDS – no. | - | - | 4 / 8 |
| **Laboratory values** |  |  |  |
| C-reactive protein (mean ± SD) | 1.21 ± 1.61 | 29.87 ± 51.60* | 89.46 ± 81.23** |
| Lactate dehydrogenase (% > upper limit of normal) | 0% | 28% | 73.53 %* |
| Hemoglobin (mean ± SD, [g/l]) | 139.88 ± 13.31 | 132.36 ± 28.33 | 128.18 ± 24.62 |
| Absolute platelet count (mean ± SD, [G/l]) | 257.25 ± 60.50 | 203 ± 67.97 | 219.74 ± 117.14 |
| Total white blood cell count (mean ± SD, [G/l]) | 5.74 ± 1.52 | 5.83 ± 2.90 | 6.65 ± 3.49 |
| Monocytes (mean ± SD, [G/l]) | 0.42 ± 0.15 | 0.50 ± 0.36 | 0.45 ± 0.34 |
| Neutrophils (mean ± SD, [G/l]) | 3.17 ± 1.03 | 3.74 ± 2.70 | 5.28 ± 3.36** |
| Eosinophils (mean ± SD, [G/l]) | 0.13 ± 0.08 | 0.04 ± 0.05* | 0.03 ± 0.07** |
| Basophils (mean ± SD, [G/l]) | 0.04 ± 0.02 | 0.02 ± 0.03* | 0.01 ± 0.02* |
| Lymphocytes (mean ± SD, [G/l]) | 1.95 ± 0.74 | 1.50 ± 0.69* | 0.81 ± 0.44** |
| CD3- CD56<sup>bright</sup> CD16<sup>dim</sup> NK cells (mean ± SD, [cells/ul]) | 11.14 ± 5.65 | 8.19 ± 4.98 | 6 ± 4.39* |
| CD3- CD56<sup>dim</sup> CD16<sup>bright</sup> NK cells (mean ± SD, [cells/ul]) | 206.29 ± 107.13 | 165.96 ± 147.96 | 162.16 ± 103.69 |
| **Level of care at blood sampling timepoint** |  |  |  |
| Outpatient – no. (%) | - | 14 (50) | - |
| Hospitalized – no. (%) | - | 14 (50) | 38 (100) |
| **Comorbidities** |  |  |  |
| Hypertension – no. (%) | - | 7 (25) | 22 (57.9)* |
| Diabetes – no. (%) | 1 (4.5) | 4 (14.3) | 12 (31.6) |
| Heart disease – no. (%) | - | 3 (10.7) | 17 (44.7)* |
| Cerebrovascular disease – no. (%) | - | 1 (3.6) | 4 (10.5) |
| Lung disease – no. (%) | - | 3 (10.7) | 6 (15.8) |
| Kidney disease – no. (%) | - | 7 (25) | 10 (26.3) |
| Malignancy – no. (%) | - | - | 4 (10.5) |
| Systemic Immunosuppression – no. (%) | - | 3 (10.7) | 4 (10.5) |
| * Indicates significance (p-value threshold <0.05) compared to the healthy, ** in the severe indicates significance in comparison to the healthy and the mild.<br><sup>a</sup> IQR denotes the interquartile range<br><sup>b</sup> COVID-19 severity at sampling according to WHO guidelines (World Health Organization, 2020) |  |  |  |

**Figure 1.**
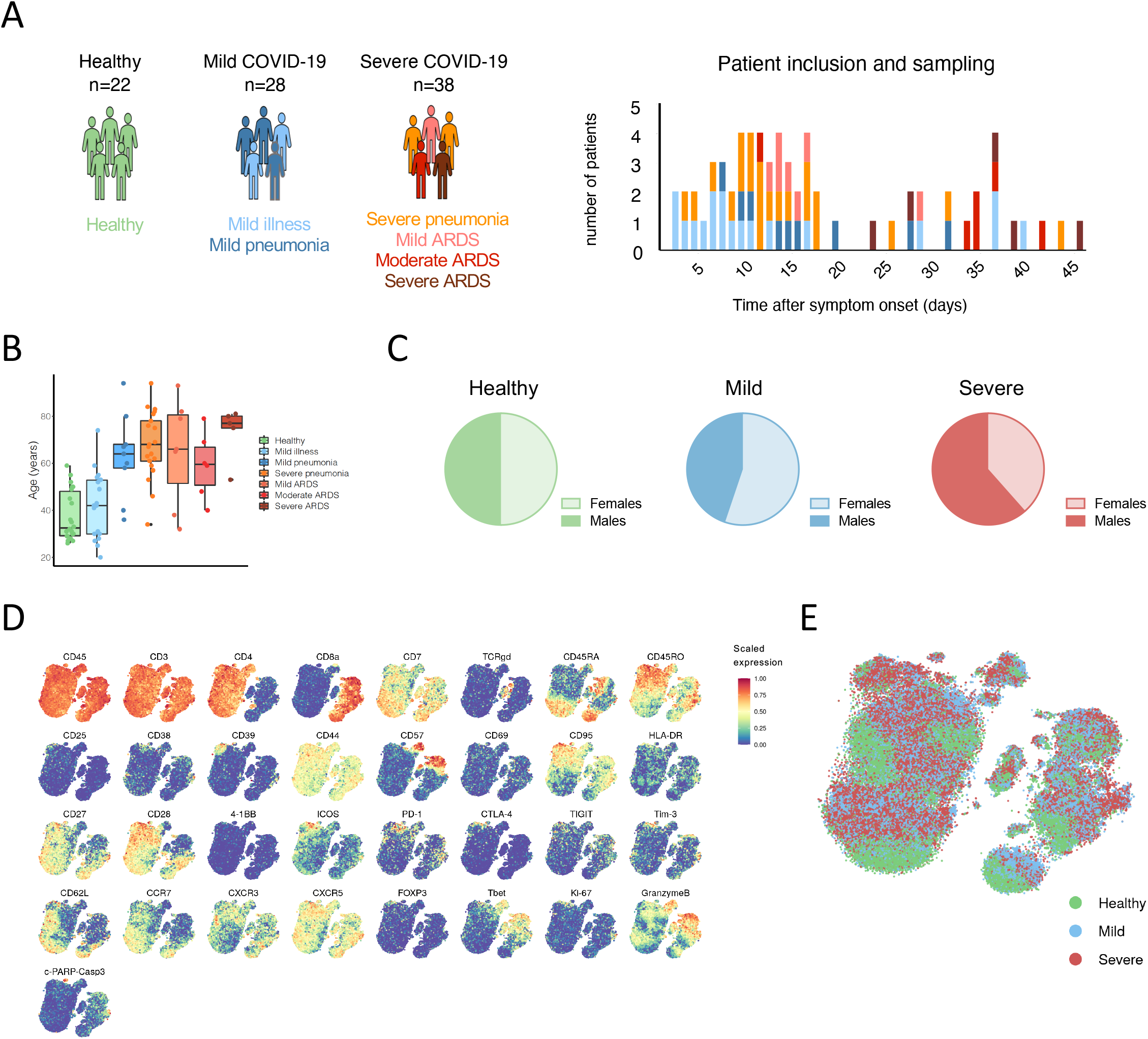
Characteristics of COVID-19 patients and healthy subjects included in the study. (A) Number of subjects recruited into the study (left) and time since onset of symptoms (right). (B) Age distribution of controls and of patients by disease severity subcategories. (C) Gender distribution of healthy subjects, patients with mild disease, and patients with severe disease. (D) t-SNE plots of normalized marker expression for 1000 T cells from all samples analyzed by mass cytometry. (E) t-SNE plot of the T cells of our study colored by disease severity.

We performed a comprehensive T cell characterization by staining PBMCs with 41 antibodies (**Table S1**) and analyzing cells using mass cytometry. T cell-related markers were visualized on t-distributed Stochastic Neighbor Embedding (t-SNE) maps (**Fig. 1d**) (Van Der Maaten and Hinton, 2008). By coloring the t-SNE map by disease severity, we observed notable differences between healthy donors and patients with mild or severe COVID-19 (**Fig. 1e)**.

### A decrease in CD8^+^ T cell subsets is a hallmark of severe COVID-19

We determined the frequencies of naive, central memory, effector memory, and terminal effector memory expressing CD45RA (TEMRA) CD4^+^ and CD8^+^ T cell subsets among all T cells in COVID-19 patients and controls (**Fig. 2a, 2b**). As previously reported (Mathew *et al.*, 2020), patients with severe disease had decreased frequencies of naive cells within the CD4^+^ and CD8^+^ T cell compartments compared to controls (**Fig. 2a-d**). The frequencies of CD4^+^ memory subsets were relatively stable (**Fig. 2c**). Conversely, the percentages of central memory CD8^+^ T cells and TEMRA CD8^+^ T cells were increased in symptomatic COVID-19 patients (**Fig. 2d**).

**Figure 2.**
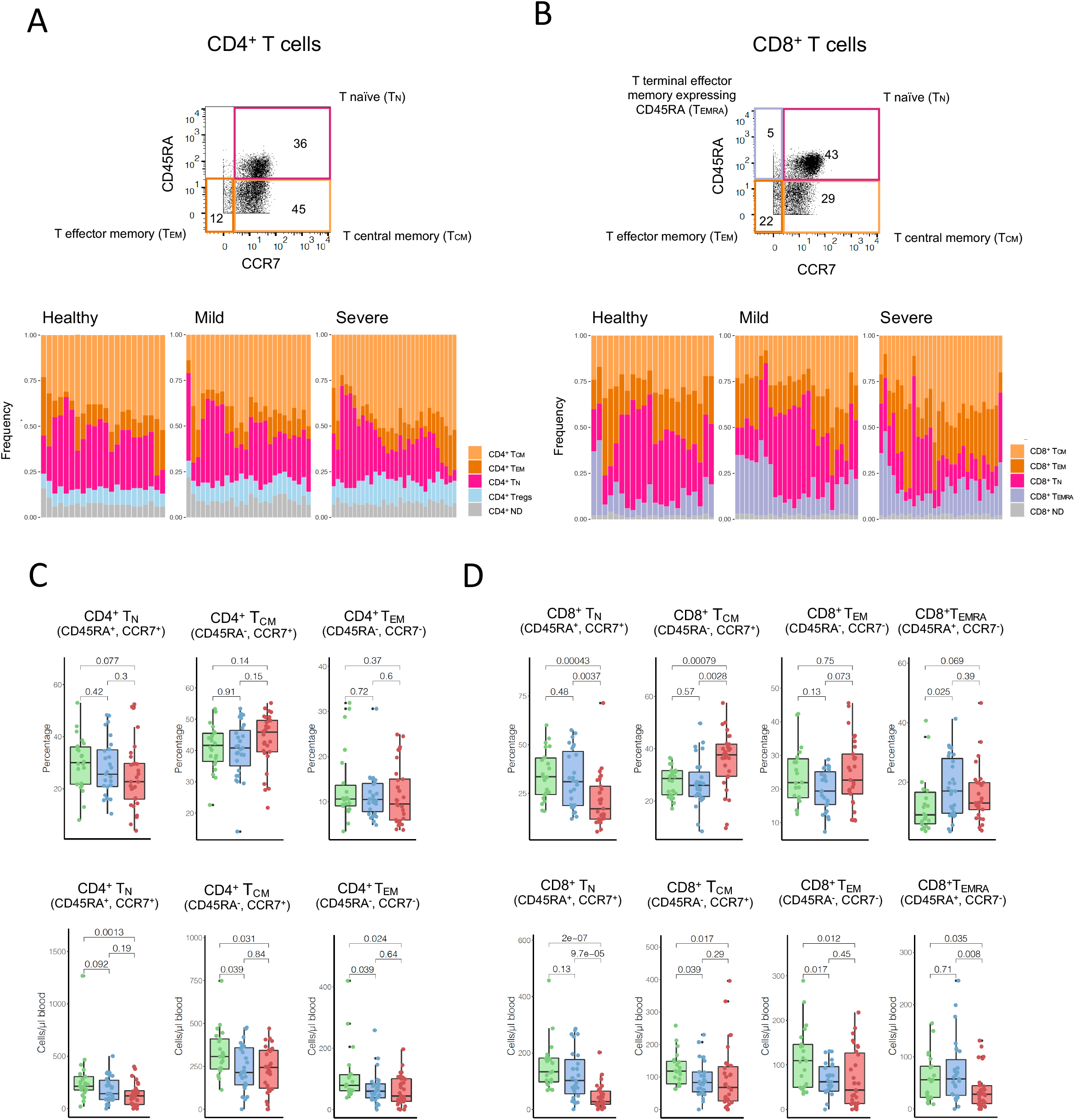
Pronounced reduction of naïve T cells during severe COVID-19 disease is not accompanied by peripheral memory expansion. (A) Gating strategy for naive and memory populations as shown on representative plots for CD4^+^ and CD8^+^ T cells (top) and stacked histograms with frequencies of regulatory T cells and naive and memory (central memory, effector memory) CD4^+^ T cells for healthy controls and mild and severe disease categories (bottom). (C) Percentages (top) and absolute counts (bottom) of CD4^+^ T cell subsets in healthy subjects and patients with mild and severe COVID-19 shown as median and interquartile ranges. Indicated p values were calculated with a Mann-Whitney-Wilcoxon test and adjusted for multiple comparisons with the Holm method. (D) Percentages (top) and absolute counts (bottom) of CD8^+^ T cell subsets in healthy subjects and patients with mild and severe COVID-19 shown as median and interquartile ranges. Statistical testing was performed as in (C).

In order to investigate how the relative changes observed within the different T cell subsets of our COVID-19 cohort related to absolute cell counts, we performed flow cytometry of whole blood samples from each subject. This enabled us to measure absolute cell counts for CD4^+^ and CD8^+^ T cells (**Fig. S2a**), which facilitated the calculation of absolute numbers for the different T cell subpopulations within our mass cytometry data set. The marked reduction in naive CD4^+^ and CD8^+^ T cells was confirmed by absolute counts and was especially pronounced for naive CD8^+^ T cells (**Fig. 2c, 2d**). In contrast, central memory and effector memory populations within the CD4^+^ T cell compartment showed a modest reduction in COVID-19 patients compared to healthy controls (**Fig. 2c**). Of note, regulatory T cells were not present in higher numbers in absolute counts in patients compared to controls, despite considerable expansion in terms of percentages (**Fig. S2b**).

Absolute numbers of central memory CD8^+^ T cells, effector memory CD8^+^ T cells, and TEMRA CD8^+^ T cells were all reduced in patients with severe disease compared to controls and to patients with mild disease (**Fig. 2e**). Thus, in severe COVID-19, naive T cell reduction was not accompanied by memory T cell expansion but rather by a slight decrease of the memory compartment, especially among CD8^+^, resulting in a decrease in peripheral T cell counts across all subsets. The patients with severe COVID-19 in our cohort were older than patients with mild disease (**Fig. 1b**), which could account for some differences in the distribution of T cell populations as previously proposed (Chen, Kelley and Goldstein, 2020). Generally, both lymphopenia and naive T cell reduction correlated with age (**Fig. S2c**). Since age and clinical severity are strongly linked, it is conceivable that some of the alterations within the T cell compartment might be due to immunological aging and thus pre-date COVID-19. However, upon stratification of patients according to age, we detected differences between mild and severe COVID-19 patients in terms of total T cell counts and naive T cell counts (**Fig. S2d**), which indicates that age alone does not explain the observed changes.

### Increased apoptosis and migration contribute to lymphopenia in severe COVID-19

We assessed different mechanisms that might contribute to peripheral T cell loss in patients with severe COVID-19. As expected, COVID-19 patients had strong T cell activation across all the main subsets (**Fig. S3a**) as well as an increase in the proportion of activated T cells and terminally differentiated and senescent T cells (**Fig. S3b, S3c**). This activation state was accompanied by considerable expansion of exhausted T cells, especially in patients with severe disease (**Fig. S4a**), and these features were present even in patients who were sampled early in the disease course (**Fig. S4b**).

Given the extent of T cell activation and exhaustion in peripheral blood, we wondered whether the T cell reduction in this compartment could be the result of activation-induced T cell death, or could rather be explained by migration of activated T cells into inflamed tissues.

Excessive proinflammatory cytokine signaling, especially mediated by TNF-α, can lead to T cell apoptosis (Mehta, Gracias and Croft, 2018). As TNF-α levels were considerably elevated in our cohort (**Fig. S4c**) in agreement with previous reports (Vabret *et al.*, 2020), we hypothesized that apoptosis could drive T cell loss during severe COVID-19 disease. When we examined the percentage of apoptotic cells among the main CD4^+^ and CD8^+^ T cell subsets, we indeed observed significant increases in the percentages of apoptotic cells among central and effector memory CD4^+^ T cells, as well as naive, central memory, and effector memory CD8^+^ T cells and TEMRA CD8^+^ T cells (**Fig. 3a**).

**Figure 3.**
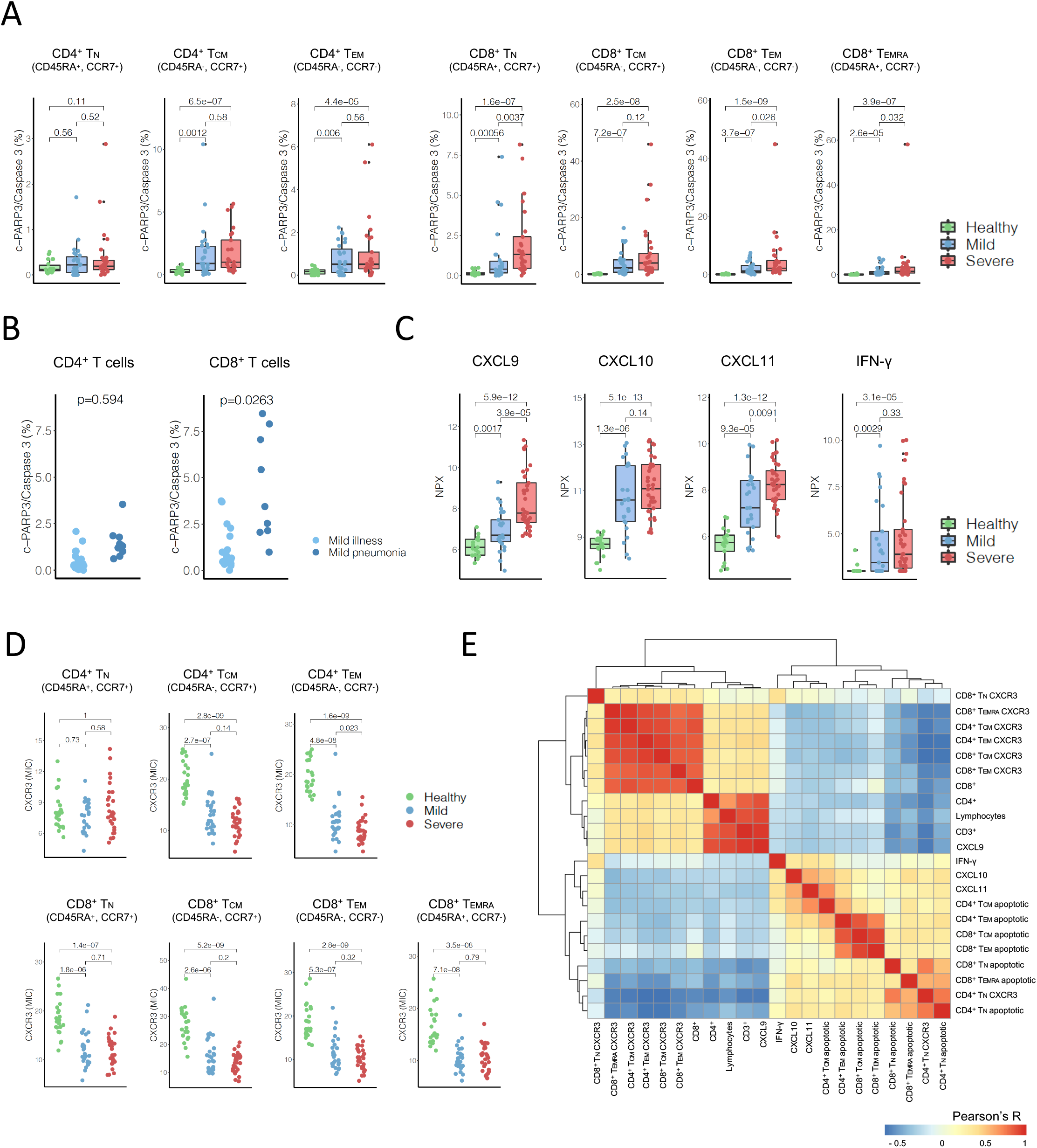
Migration and apoptosis contribute to lymphopenia in patients with severe COVID-19. (A) Percentages of apoptotic (cleaved-PARP/cleaved Caspase 3^+^) cells among CD4^+^ T cell subsets and CD8^+^ T cell subsets in healthy subjects and patients with mild and severe COVID-19 patients shown as median and interquartile ranges. Indicated p values were calculated with a Mann-Whitney-Wilcoxon test and adjusted for multiple comparisons with the Holm method. (B) Percentages of apoptotic (cleaved-PARP/cleaved Caspase 3^+^) cells among CD4^+^ T cells and CD8^+^ T cell subsets in patients with mild illness versus mild pneumonia. Indicated p values were calculated with a Mann-Whitney-Wilcoxon test. (C) CXCL9, CXCL10, CXCL11, and IFN-γ serum levels in healthy subjects and patients with mild and severe COVID-19 measured with an Olink proximity extension assay shown as median and interquartile ranges. Statistical testing was performed as in (A). (D) CXCR3 mean ion count (MIC) of CD4^+^ T cell subsets (top) and CD8^+^ T cell subsets (bottom) in healthy subjects and patients with mild and severe COVID-19. Statistical testing was performed as in (A). (E) Pearson correlation analyses among CXCR3, CXCL9, CXCL10, CXCL11, and IFN-γ levels, lymphocyte and T cell counts, and percentage of apoptotic cells in samples from all subjects.

Among CD8^+^ T cells, the extent of the apoptosis was higher in samples from patients with severe COVID-19 disease compared to those from patients with mild symptoms (**Fig. 3a**). Even within mild cases, the extent of apoptosis was strongly associated with disease severity in the CD8 subset (**Fig. 3b**). Increased T cell apoptosis could thus contribute to the lymphopenia that is a hallmark of severe COVID-19. Further supporting this hypothesis, we observed an inverse correlation between elevated apoptosis and absolute T cell numbers (**Fig. 3e**).

Next, we examined T cell migration to inflamed tissues as a possible mechanism driving peripheral lymphopenia. Lymphocyte accumulation in infected tissues such as the lung during COVID-19 progression has been reported in some studies (Liao *et al.*, 2020)(Xu *et al.*, 2020) but not others (Tian *et al.*, 2020). We therefore sought to detect indirect signs of T cell migration in the peripheral blood of COVID-19 patients. T cell migration into inflamed lung parenchyma is primarily mediated by CXCR3 signaling (Fadel *et al.*, 2008)(Kohlmeier *et al.*, 2009), and we therefore investigated the level of CXCR3 ligands in the serum of COVID-19 patients. CXCR3 ligands CXCL9, CXCL10, and CXCL11 were significantly increased in sera from COVID-19 patients, especially in sera from patients with severe disease (**Fig. 3c**). Furthermore, IFN-γ is a potent inducer of CXCR3 ligands (Van Raemdonck *et al.*, 2015), and we observed a marked elevation in IFN-γ (**Fig. 3c and Fig. S4d**). We also observed a very strong reduction in CXCR3 expression in all the main T cell subsets with the exception of naive CD4^+^ T cells in symptomatic patients (**Fig. 3d and S4e**). Furthermore, CXCR3 abundance inversely correlated with levels concentrations of CXCL9, CXCL10, CXCL11 and IFN-γ, and was positively correlated with levels of lymphocytes, CD3^+^ T cells, CD4^+^ T cells, and CD8^+^ T cells levels (**Fig. 3e**). These data are suggestive of a role for T cell migration in the lymphopenia associated with severe COVID-19.

### Patients with severe COVID-19 show signs of IL-7-induced homeostatic proliferation of T cells

Because migration and apoptosis are both likely to contribute to the observed T cell lymphopenia, we looked for indirect signs that would indicate a global T cell loss versus a shift from the blood compartment to the inflamed tissues.

IL-7 is known to be a critical homeostatic factor for T cells and, since IL-7 production by stromal cells is relatively constant (Surh and Sprent, 2008), its levels are mostly regulated by lymphocyte consumption. In patients with severe disease, we observed higher serum IL-7 levels compared to those with mild disease (**Fig. 4a**). IL-7 elevation was independent of time since symptom onset (**Fig. 4b**). Furthermore, IL-7 levels inversely correlated with the total number of CD8^+^ T cells and there was a tendency toward inverse correlations with CD4^+^ T cell counts (**Fig. 4c**). Taken together, these findings indicate a global T cell loss in severe COVID-19, which results in a systemic IL-7 elevation.

**Figure 4.**
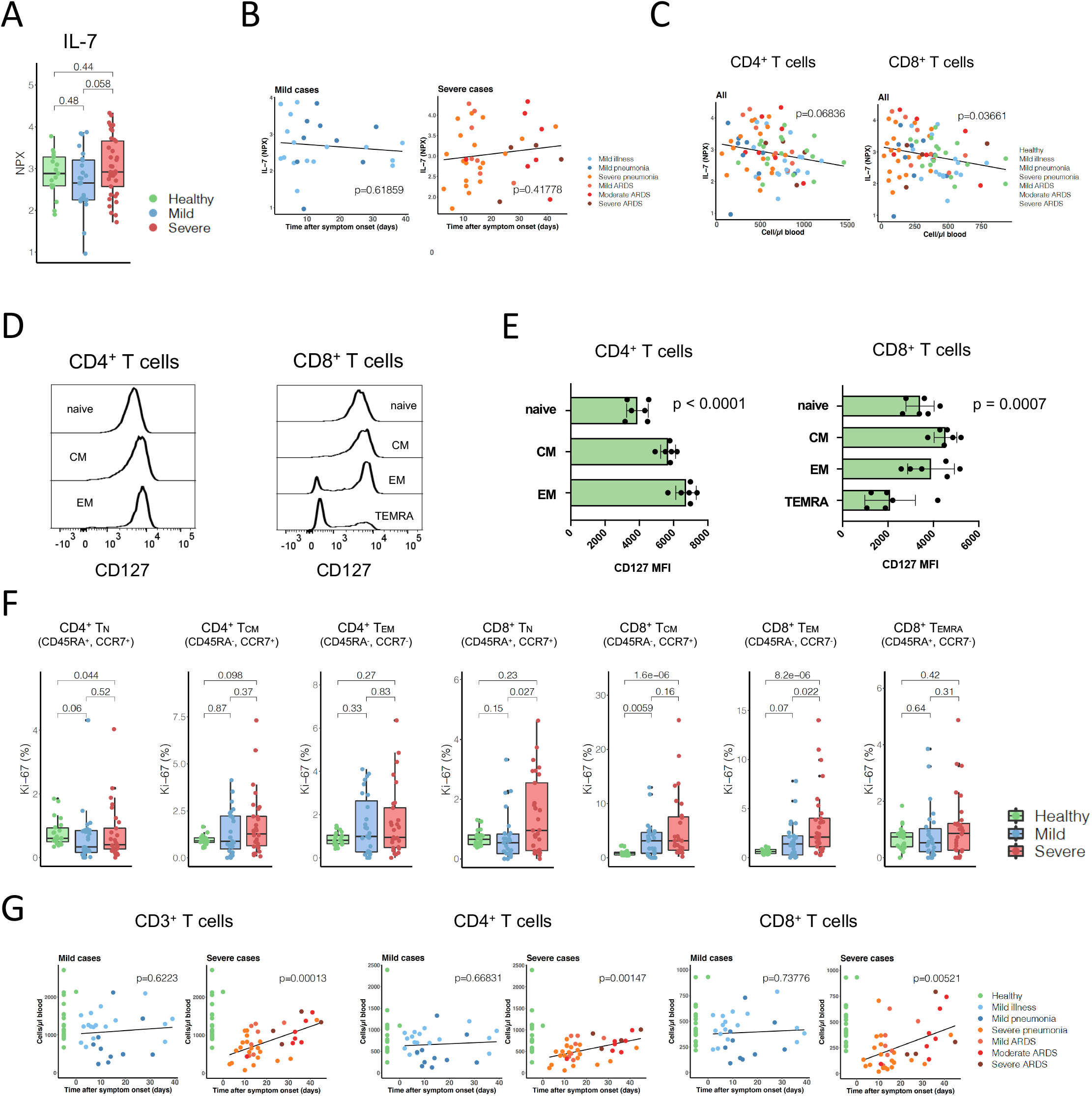
Patients with severe COVID-19 show signs of IL-7-induced homeostatic proliferation. (A) IL-7 serum levels in healthy subjects and patients with mild and severe COVID-19 measured with the Olink proximity extension assay shown as median and interquartile range. Indicated p values were calculated with a Mann-Whitney-Wilcoxon test and adjusted for multiple comparisons with the Holm method. (B) IL-7 serum levels are a function of time after symptom onset in mild and severe COVID-19 patients as shown by a linear model. (C) IL-7 serum level is a function of CD4^+^ T cells, and CD8^+^ T cells as shown by a linear model. (D) Representative plots of CD127 expression CD4^+^ T cell and CD8^+^ T cell subsets in healthy subjects. Indicated p values were calculated with a Kruskal-Wallis test. (E) CD127 mean fluorescence intensity (MFI) of CD4^+^ T cell and CD8^+^ T cell subsets in healthy subjects. (F) Percentages of proliferating (Ki-67^+^) cells among CD4^+^ T cell subsets (top) and CD8^+^ T cell subsets (bottom) in healthy subjects and patients with mild and severe COVID-19 shown as median and interquartile ranges. Indicated p values were calculated with a Mann-Whitney-Wilcoxon test and adjusted for multiple comparisons with the Holm method. (G) CD3^+^, CD4^+^, and CD8^+^ T cell counts are a function of time after symptom onset in patients with mild and severe COVID-19 as shown by a linear model. Counts for healthy subjects are shown for reference but were not included in the model.

To assess which T cells would be most responsive to increased IL-7, we determined protein abundance of IL-7 receptor α (also termed CD127) on naive and memory T cell subsets. We observed lower abundance of IL-7 receptor α on TEMRA cells compared to naive, central memory, and effector memory CD4^+^ and CD8^+^ T cells of healthy subjects (**Fig. 4d** and **4e**). When we looked at T cell proliferation in those same compartments in COVID-19 patients, we did not observe changes in the percentage of proliferating (Ki-67^+^) cells as compared to healthy controls in the CD4^+^ compartment (**Fig. 4f**). In contrast, we observed higher frequencies of proliferative naive, central memory and effector memory CD8^+^ T cells in COVID-19 patients. Proliferation was especially prominent in patients with severe disease. Of note, we did not observe increased proliferation of TEMRA CD8^+^ T cells, despite their activation (**Fig. S3a**), in agreement with their reduced IL-7 sensitivity.

Perhaps as a result of proliferation, we observed higher T cell counts in samples from patients with severe COVID-19 taken at later time points after symptom onset than in samples taken earlier in the disease course (**Fig. 4g**).

### Lymphopenia induced T cell proliferation drives expansion of poly-specific T cells

We next investigated whether the observed early lymphopenia and subsequent proliferation had an impact on T cell function. We assessed the ability of peripheral T cells from patients with mild and severe COVID-19 to form blasts in response to mitogens, superantigens, and a selection of common viral antigens. We observed markedly less blast formation by PBMCs from patients upon stimulation with Varicella Zoster Virus (VZV), Herpes simplex virus 1 (HSV1), Herpes simplex virus 2 (HSV2) and adenovirus as compared to healthy controls, and the difference was more pronounced in samples from the severe COVID-19 group than from the mild groups (**Fig. 5a**). The same tendency was not observed when PBMCs were incubated with pokeweed mitogen or *Staphylococcus* superantigen, but we did observe a reduction in proliferation upon stimulation with concanavalin A (**Fig. S5a**).

**Figure 5.**
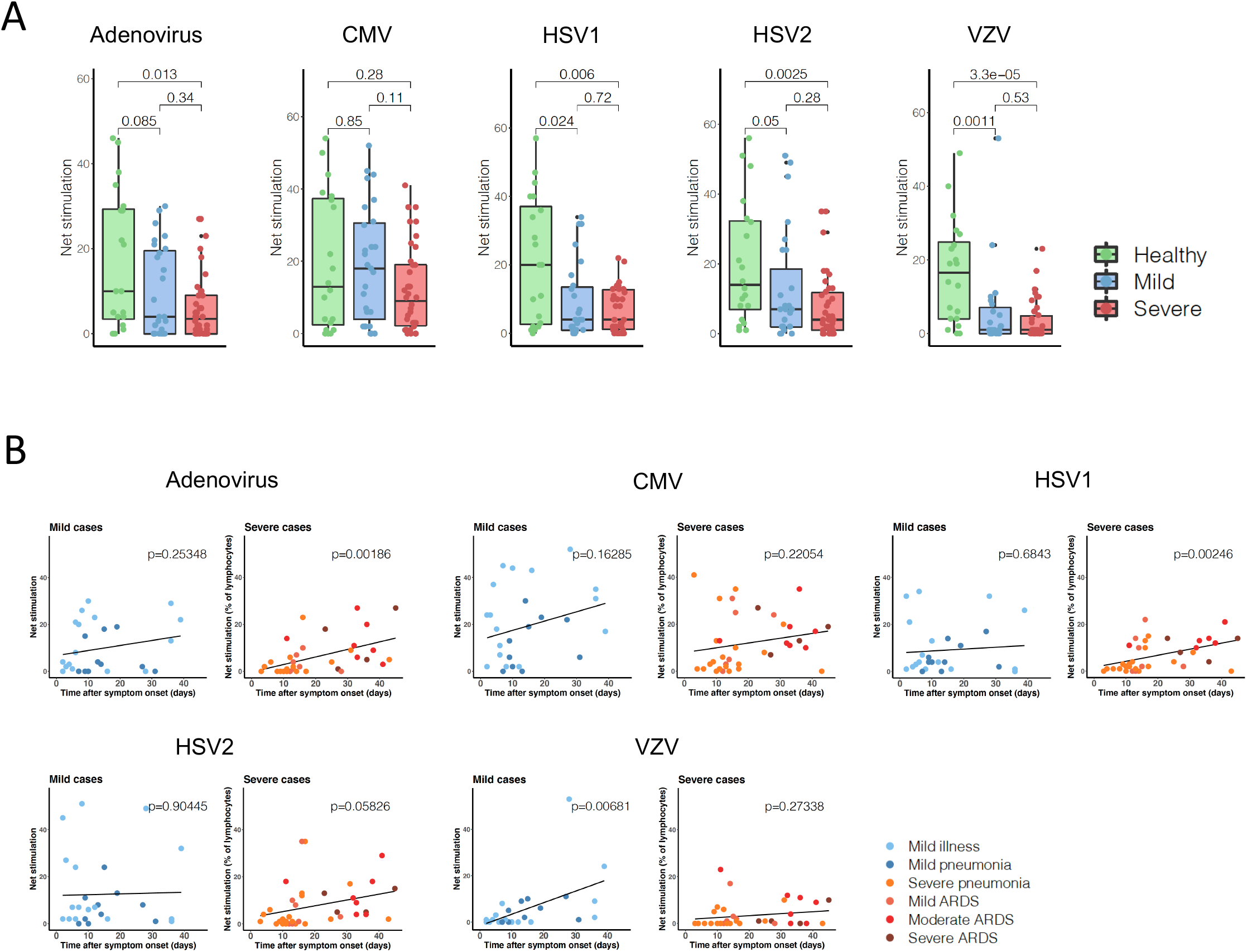
Homeostatic proliferation drives expansion of poly-specific T cells. (A) Net stimulation of CD3^+^ T cells observed in the presence of viral antigens shown as median and interquartile ranges. Indicated p values were calculated with a Mann-Whitney-Wilcoxon test and adjusted for multiple comparisons with the Holm method. (B) Linear modeling of blast formation as a function of time after symptom onset in mild and severe COVID-19.

Although reduced blast formation upon virus stimulation is likely determined by a combination of factors, reduced frequency of precursors of virus-specific memory T cells could be an important element. In support of this, we found that blast formation positively correlated with CD4^+^ and CD8^+^ T cell counts (**Fig. S5b, S5c**). Oligoclonal expansion of SARS-CoV-2-specific T cell clones could account for reduced peripheral precursor memory T cell frequency in mild COVID-19 patients; these patients did not have significant lymphopenia but did have diminished functional T cell responses.

To determine whether lymphopenia induced homeostatic proliferation could contribute to the observed T cell proliferation, especially in severe COVID-19, we used the different sampling time of our included patients as a pseudo-longitudinal time course. Since blast formation in our functional assay reflects – at least to some extent – specific memory T cell precursor frequency, we reasoned that homeostatic proliferation would lead to an increase in poly-specific memory T cell over time, which in turn would lead to increased blast formation upon stimulation with specific viral antigens.

Indeed, when we evaluated the relationship between blast formation in response to different viral stimuli and the time between symptom onset and sample collection, we detected a marked increase in blast formation upon stimulation with HSV1, HSV2, and adenovirus as a function of time of sample collection after symptom onset (**Fig. 5b**). A trend toward increased blast formation with time was also observed for CMV and VZV stimulation of samples from patients with severe disease (**Fig. 5b**), but this was not observed upon nonspecific stimulation with mitogens and superantigens (**Fig. S5d**). The increased blast formation at later time points was also positively correlated with absolute T cell counts (**Fig. S5b, S5c**) further linking lymphopenia, homeostatic T cell proliferation, and increased poly-specific functional capacity. In summary, in patients with severe COVID-19, lymphopenia induced homeostatic proliferation is present and likely drives expansion of poly-specific T cells, leading to an improvement of T cell functionality over time.

### Unbiased clustering of the T cell compartment confirms that T cell activation, exhaustion, and apoptosis are associated with disease severity

In order to investigate whether phenotypes identified in an unbiased manner across all T cell populations would show the same trends as our analysis of pre-defined cell subsets, we performed automated clustering using the PhenoGraph algorithm. Automated clustering identified 15 phenotypically distinct T cell clusters. Clusters were manually annotated based on the most prominent signature of marker expression (**Fig. 6a**). Marked differences in the frequencies of the identified clusters were evident in healthy donors, patients with mild COVID-19, and patients with severe COVID-19 (**Fig. 6b**). In agreement with our analysis of pre-defined subsets, we observed a marked reduction in naive CD8^+^ T cells in patients with severe disease, whereas the proportion of naive CD4^+^ T cells remained stable (**Fig. 6c**). Frequencies of both the activated T cell subsets (clusters 9 and 10) and the apoptotic, exhausted subsets (clusters 12 and 15) were markedly increased in patients with severe disease compared to controls (**Fig. 6c** and **Fig. S6**). We also observed expansion of granzyme B-producing cytotoxic CD4^+^ T cells in patients, especially those with severe COVID-19. Frequencies of some clusters were strongly associated. This was the case for TEMRA CD8^+^ cells and apoptotic or exhausted CD8^+^ T cells, activated and regulatory CD4^+^ T cells, as well as apoptotic exhausted CD4^+^ T cells (**Fig. 6d**). In summary, an unbiased analysis of the mass cytometry data confirmed our key findings on naïve T cell reduction and substantial increase in T cell activation, exhaustion and apoptosis.

**Figure 6.**
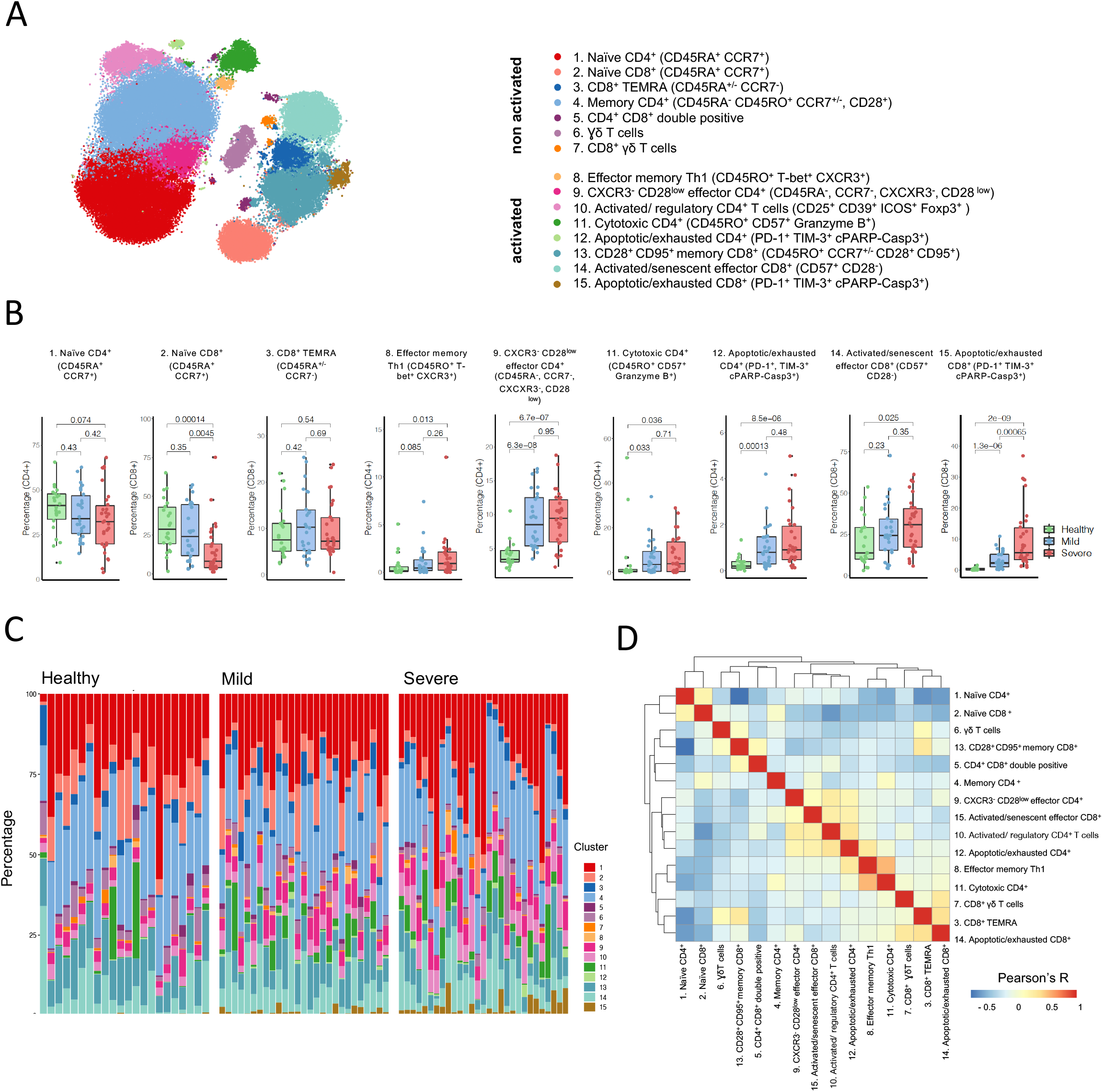
T cell activation, exhaustion, and apoptosis occurs during COVID-19 progression. (A) t-SNE of T cells colored by PhenoGraph clusters. 1000 cells are shown for each sample. (B) Frequencies of PhenoGraph T cell clusters calculated among CD4+ and CD8+ T cells in healthy subjects and patients with mild and severe COVID-19 shown as median and interquartile ranges. Indicated p values were calculated with a Mann-Whitney-Wilcoxon test and adjusted for multiple comparisons with the Holm method. (C) Frequencies of PhenoGraph T cell clusters in healthy subjects and patients with mild and severe COVID-19. (D) Pearson correlation analysis using the frequencies of the PhenoGraph T cell clusters.

## Discussion

Lymphopenia and hyperinflammation are very prominent features of severe COVID-19, which speaks to the importance of immunological processes for the disease pathogenesis. Since T cells are central players in the control of viral infections, understanding of the perturbances of T cell homeostasis in COVID-19 can provide insight into the disease mechanisms and inform therapeutic strategies. In our well characterized COVID-19 cohort, we were able to dissect changes in the T cell compartments and relate them to disease duration, which allowed us to appreciate dynamic changes in T cell populations. We were able to identify both extensive apoptosis and migration as possible mechanisms driving the lymphopenia, as well as global increase in T cell counts over a pseudo-longitudinal time course, which was at least partially due to homeostatic proliferation.

The occurrence of peripheral lymphopenia has been described in several human acute respiratory viral infections (Lewis, Gilbert and Knight, 1986)(Russell *et al.*, 2017). Lymphopenia in this setting is thought to be in part caused by trafficking of activated lymphocytes into inflamed tissues where viral replication is ongoing (Kohlmeier *et al.*, 2009). In severe COVID-19, the extent of lymphopenia is closely linked to disease severity, development of ARDS, and mortality, which suggests a central role for T cell lymphopenia in COVID-19 immunopathology (Vabret *et al.*, 2020). Since disease severity in COVID-19 is associated with biological age, age-associated decline in lymphocytes could in part account for the observed lymphopenia. Of note, the main age-related alteration of the T cell compartment that has been associated with increased risk of mortality is CD4^+^ T cell loss and inversion of the CD4 to CD8 T cell ratio (Hadrup *et al.*, 2006). In severe COVID-19, however, we and others have shown an extensive preferential loss of peripheral CD8^+^ T cells (Mathew *et al.*, 2020), which suggests that other mechanisms than age related changes may be involved.

Our analysis of absolute T cell counts indicated marked T cell loss across all subsets in the CD4^+^ and CD8^+^ T cell compartment in patients with COVID-19 compared to healthy controls. Thus, specific mechanisms directly related to SARS-CoV-2 infection appear to contribute to the observed changes. Our finding of pronounced elevation in the levels of the CXCR3 ligands CXCL9, CXCL10, and CXCL11 in conjunction with reduced surface expression of CXCR3 on several key T cell subsets in patients with symptomatic COVID-19 suggests that migration of T cells into inflamed tissues contributes to the observed lymphopenia. However, the comparable reduction in CXCR3 expression on both CD4^+^ and CD8^+^ T cell subsets does not account for the observed preferential loss of CD8 T cells. In support of our hypothesis that increased T cell migration into inflamed tissues does not fully explain the observed reduction in peripheral T cell numbers, we found evidence of increased amounts of circulating IL-7 in patients with severe COVID-19. Levels of circulating IL-7 are primarily regulated by its consumption by lymphocytes (Surh and Sprent, 2008); thus, the observed reduced peripheral T cell numbers suggests a global T cell loss in addition to lymphocyte redistribution in the setting of an acute viral infection. We also observed that the frequency of apoptotic CD8^+^ T cell subsets in patients with severe COVID-19 was far greater than among CD4^+^ T cells indicating that apoptosis of CD8^+^ T cells could explain the preferential loss of this subset and be a central mechanism in the immunopathology of severe COVID-19. Whether T cell apoptosis occurs as a result of activation-induced cell death in the setting of extensive inflammation or upon direct interaction with the SARS-CoV-2 virus remains to be elucidated.

We did not observe changes in the percentage of proliferating CD4 T cells in our COVID-19 patients, but there was evidence for increased frequencies of proliferative, naïve, central memory and effector memory CD8^+^ T cells that was especially prominent in patients with severe disease. Although proliferation in the CD8^+^ compartment could be due to expansion of SARS-CoV-2-specific T cells, the increased levels of IL-7 and the robust proliferation of naive CD8^+^ T cells, especially in severe disease, suggest a role for IL-7-induced homeostatic proliferation. Furthermore, in patients with severe COVID-19 we observed an increase in blast formation upon stimulation with common viral antigens in our pseudo-longitudinal time course. Since blast formation correlated well with overall CD4^+^ and CD8^+^ T cell counts, we interpret this as evidence of increased precursor frequency at later time points during the infection. Taken together, these data indicate that homeostatic expansion of poly-specific memory T cells occurs in COVID-19 patients with severe disease.

In our cohort, we also found evidence of increased T cell exhaustion soon after the onset of symptoms as well as impaired T cell function, as evidenced by reduced blast formation upon stimulation with viral antigens. Early T cell dysfunction in patients who eventually develop severe disease might be caused by an insufficient initial type I interferon response as recently suggested (Hadjadj *et al.*, 2020). The ensuing suboptimal T cell immunity could then facilitate further viral dissemination that in turn stimulates innate immunity, perhaps through CD16^+^ monocytes (Chevrier *et al.,* 2020). The subsequent increase in production of pro-inflammatory cytokines could culminate in a cytokine storm and development of ARDS.

To what extent age-associated individual immunological predisposition contributes to the development of severe COVID-19 could not be determined in the current analysis. Further longitudinal immunological studies of patients recovering from COVID-19 will likely be able to clarify this important aspect. In addition to IL-7, IL-15 is also important in driving lymphopenia-induced homeostatic proliferation, particularly of memory CD8^+^ T cells (Raeber *et al.*, 2018). IL-15 is usually bound to the IL-15 receptor α (Dubois *et al.*, 2002)(Mortier *et al.*, 2008)(Bouchaud *et al.*, 2013), which means that serum concentrations of soluble IL-15 are an unreliable measurement of the availability of bioactive IL-15. However, IL-15 elevation in patients with COVID-19 has been reported (Hadjadj *et al.*, 2020) and could likely contribute to the increase we observed in CD8^+^ T cell proliferation. We did not investigate SARS-CoV-2-specific T cells in the current study. Determining the frequency of virus-specific cells in each T cell compartment, and especially in the proliferating fraction, will shed light on the mechanisms of T cell proliferation and will assist in defining the extent of homeostatic proliferation.

Understanding the immunological mechanisms underlying severe COVID-19 disease is necessary for risk stratification and central to the development of interventional therapies as well as protective vaccines. Our study reveals that severe COVID-19 is associated with extensive loss of T cells, especially within the CD8^+^ T cell compartment. Since the observed T cell apoptosis is closely associated with the destructive inflammatory environment of severe COVID-19 our study highlights the potential of anti-inflammatory therapies in possibly preventing the observed extensive T cell loss.

## Author contribution

SA, and SC contributed to study design, patient recruitment, data collection, data analysis and data interpretation. CC and YZ contributed to patient recruitment, data collection and data analysis. MER, EB, AR, MS-H, LCH, and DJS contributed to patient recruitment and clinical management. SS, AJ and SC developed the Cytof antibody panel and performed the CyTOF experiments. SA and JN wrote the manuscript with assistance from OB and BB. JN, OB and BB contributed to study conception and design, data analysis and data interpretation. All authors reviewed and approved the final version of the manuscript.

## Acknowledgments

We thank Alessandra Guaita, Jennifer Jörger, Sara Hasler and the members of the Boyman laboratory for their support of the study. We thank Natalie de Souza for helpful discussions.

