## Supplementary Material for "Lymphopenia-induced T cell proliferation is a hallmark of severe COVID-19"

### Supplemental Figure Legends

**Figure S1. Batch correction for mass cytometry.**

A) Schematic of the strategy used to collect mass cytometry data. Samples were analyzed in two batches. The indicated number of samples were barcoded and stained with the same antibody mix stored as frozen aliquots.

(B) t-SNE plot calculated on the three reference samples included in each CyTOF run based on all markers included in the study, colored by batch.

(C) Histograms of signal intensities for each marker included in the panel observed on the pooled references acquired in the two runs. Data were linearly scaled based on the 98$^{th}$ percentile. A small shift in background intensity was observed for Granzyme B.

**Figure S2. T cell absolute counts and age-related changes in T cell populations.**

(A) Absolute counts of lymphocytes and CD3$^{+}$, CD4$^{+}$, and CD8$^{+}$ T cells in healthy subjects and patients with mild and severe COVID-19 shown as median and interquartile ranges. Indicated p values were calculated with a Mann-Whitney-Wilcoxon test and adjusted for multiple comparisons with the Holm method.

(B) Percentages and absolute counts of regulatory T cells in healthy subjects and patients with mild and severe COVID-19 are shown as median and interquartile ranges. Statistical testing was performed as in (A).

(C) A linear model of CD4$^{+}$ and CD8$^{+}$ T cell counts and naive CD4$^{+}$ and CD8$^{+}$ T cell counts as a function of age in COVID-19 patients shows a strong relationship.

(B) Absolute counts of CD4$^{+}$ and CD8$^{+}$ T cell counts and naive CD4$^{+}$ and CD8$^{+}$ T cell counts in healthy subjects and patients with mild and severe COVID-19 younger than 60 years of age (top) and older than 60 years of age (bottom) shown as median and interquartile ranges. Statistical testing was performed as in (A).

**Figure S3. Patients with COVID-19 irrespective of disease severity have strong T cell activation.**

(A) Mean ion count (MIC) of CD25, CD69, HLA-DR, ICOS, and PD-1 on CD4$^{+}$ T cell subsets (top) and CD8$^{+}$ T cell subsets in healthy subjects and patients with mild and severe COVID-19. Indicated p values were calculated with a Kruskal-Wallis test.

(B) Percentages and absolute counts of activated (HLA-DR$^+$) CD4$^+$ and CD8$^+$ T cells in healthy subjects and patients with mild and severe COVID-19 shown as median and interquartile ranges. Indicated p values were calculated with a Mann-Whitney-Wilcoxon test and adjusted for multiple comparisons with the Holm method.
(C) Percentages and absolute counts of senescent (CD28$^-$, CD57$^+$) CD4$^+$ and CD8$^+$ T cells in healthy subjects and patients with mild and severe COVID-19 shown as median and interquartile ranges. Statistical testing was performed as in (B).

**Figure S4. T cell exhaustion, pro-inflammatory cytokine elevation and CXCR3 levels increased T cell migration in COVID-19.**

(A) Percentages and absolute counts of exhausted (PD1$^+$, TIM-3$^+$) CD4$^+$ T cells and exhausted (PD1$^+$, TIM-3$^+$) CD8$^+$ T cells in healthy subjects and patients with mild and severe COVID-19 shown as median and interquartile ranges. Indicated p values were calculated with a Mann-Whitney-Wilcoxon test and adjusted for multiple comparisons with the Holm method.
(B) Linear modeling of exhausted T cell counts as a function of time after symptom onset in patients with mild and severe COVID-19.
(c) TNF-α serum levels in healthy subjects and patients with mild and severe COVID-19 measured with a proximity extension assay (left) and ELISA (right).
Data are shown as median and interquartile ranges. Statistical testing was performed as in (A).
(d) IFN-γ serum levels in healthy subjects and patients with mild and severe COVID-19 measured with ELISA shown as median and interquartile range. Statistical testing was performed as in (A).
(E) Representative plots of CXCR3 expression on CD4$^+$ T cell subsets (top) and CD8$^+$ T cell subsets (bottom) in healthy subjects and patients with mild and severe COVID-19.

**Figure S5. Blast formation upon stimulation with viral antigens correlates with absolute T cell counts.**

(A) Net stimulation of CD3$^+$ T cells in the presence of pokeweed mitogen, concanavalin A, or *Staphylococcus* enterotoxin a and b shown as median and interquartile ranges.

Indicated p values were calculated with a Mann-Whitney-Wilcoxon test and adjusted for multiple comparisons with the Holm method.

(B) Linear modeling of blast formation upon stimulation with mitogens, superantigens, and viruses as a function of absolute $CD4^{+}$ T cell counts.

(C) Linear modeling of blast formation upon stimulation with mitogens, superantigens, and viruses as a function of absolute $CD8^{+}$ T cell counts.

(D) Linear modeling of blast formation upon stimulations with mitogens and superantigens as a function of time after symptom onset in patients with mild and severe COVID-19.

**Figure S6. Frequencies of PhenoGraph clusters.**

Frequencies of PhenoGraph T cell clusters among $CD3^{+}$, $CD4^{+}$, and $CD8^{+}$ T cells in healthy subjects and patients with mild and severe COVID-19 shown as median and interquartile ranges. Indicated p values were calculated with a Mann-Whitney-Wilcoxon test and adjusted for multiple comparisons with the Holm method.

### Table S1. Mass cytometry panel.

| Antigen | Provider | Clone | Cat No | Metal tag |
|---|---|---|---|---|
| CD15 | BioLegend | HI98 | 301902 | 89Y |
| CD57 | BioLegend | HNK-1 | 359602 | 141Pr |
| CD38 | BioLegend | HIT2 | 303502 | 142Nd |
| CD69 | eBioscience | FN50 | 14-0699-82 | 143Nd |
| CD27 | Abcam | EPR8569 | ab192336 | 144Nd |
| CD4 | BioLegend | RPA-T4 | 300502 | 145Nd |
| CD45RA | BioLegend | HI100 | 304102 | 146Nd |
| CD11b | BioLegend | M1/70 | 101202 | 147Sm |
| CD20 | BD Bioscience | H1(FB1) | 555677 | 148Nd |
| CD25 | BD Bioscience | M-A251 | 356102 | 149Sm |
| CD44 | Fluidigm | IM7 | 3150018B | 150Nd |
| CD39 | BioLegend | A1 | 328202 | 151Eu |
| FOXP3 | eBioscience | 236A/E7 | 14-4777-82 | 152Sm |
| human IgM | BioLegend | MHM-88 | 314502 | 153Eu |
| CTLA-4 | BioLegend | L3D10 | 349902 | 154Sm |
| TCR gamma/delta | BioLegend | B1 | 331202 | 155Gd |
| CXCR5 | Abcam | EPR8837 | ab225575 | 156Gd |
| human IgD | BioLegend | IA6-2 | 348202 | 158Gd |
| TIGIT | BioLegend | A15153G | 372702 | 159Tb |
| Tbet | BioLegend | 4B10 | 644801 | 160Gd |
| CD95 (Fas) | BioLegend | DX2 | 305602 | 161Dy |
| CD45RO | BioLegend | UCHL1 | 304202 | 162Dy |
| CD279 (PD-1) | BioLegend | EH12.2H7 | 329902 | 163Dy |
| CD62L | BioLegend | DREG-56 | 304802 | 164Dy |
| CD126 | BioLegend | UV4 | 352802 | 165Ho |
| CD278 (ICOS) | BioLegend | C398.4A | 313502 | 166Er |
| Tim-3 | R&D Systems | polyclonal | AF2365 | 167Er |
| CD14 | BioLegend | M5E2 | 301802 | 168Er |
| CXCR3 / CD183 | BD Bioscience | 1C6 | 557183 | 169Tm |
| 4-1BB | BioLegend | 4B4-1 | 309802 | 170Er |
| CD28 | BioLegend | 28.2 | 302902 | 171Yb |
| CCR7 | BioLegend | G043H7 | 353202 | 172Yb |
| Granzyme B | Invitrogen Antibodies | GB11 | MA1-80734 | 173Yb |
| CD11c | BioLegend | Bu15 | 337202 | 174Yb |
| CD8a | BioLegend | RPA-T8 | 301002 | 175Lu |
| cleaved PARP | BD Bioscience | F21-852 | 552596 | 176Yb |
| Cleaved Caspase3 | BD Bioscience | C92-605 | 559565 | 176Yb |
| HLA-DR | BioLegend | L243 | 307602 | 194Pt |
| CD3 | BioLegend | UCHT1 | 300402 | 195Pt |
| CD7 | BD Bioscience | M-T701 | 555359 | 196Pt |
| Ki-67 | BD Bioscience | B56 | 556003 | 198Pt |
| CD45 | BioLegend | HI30 | 304002 | 209Bi |

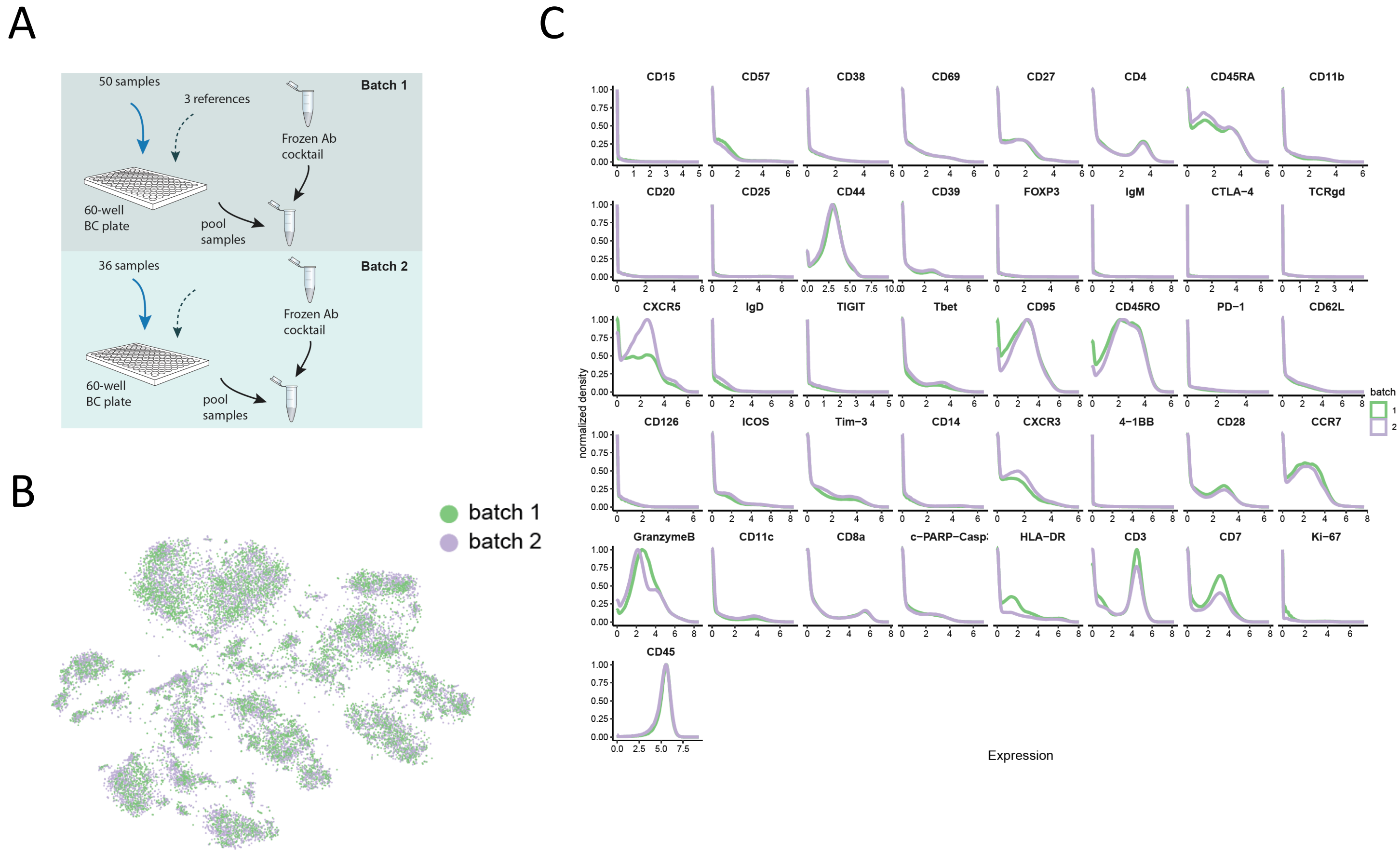


**Figure S1**. Batch correction for mass cytometry.

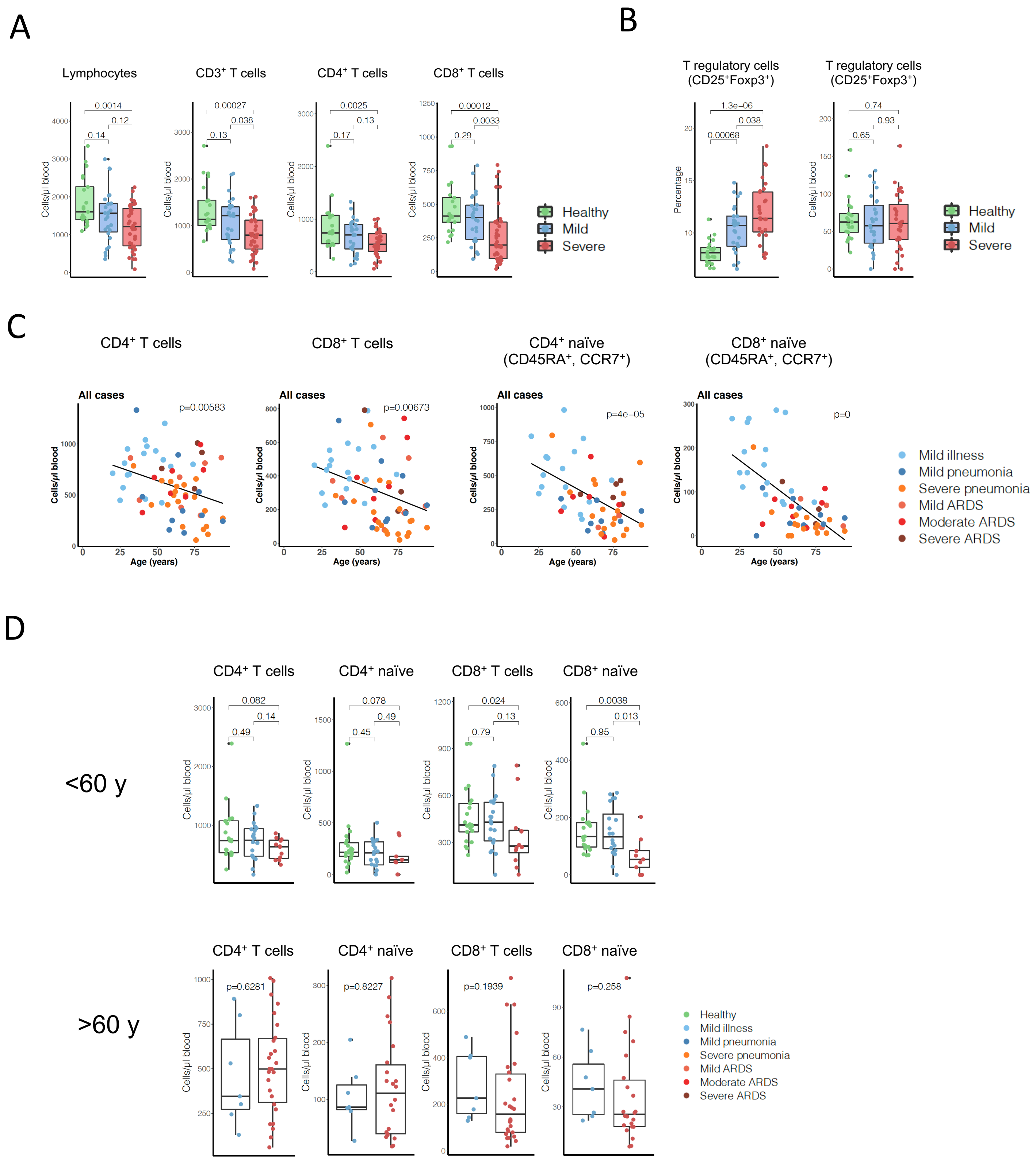


**Figure S2**. T cell absolute counts and age related changes in T cell populations.

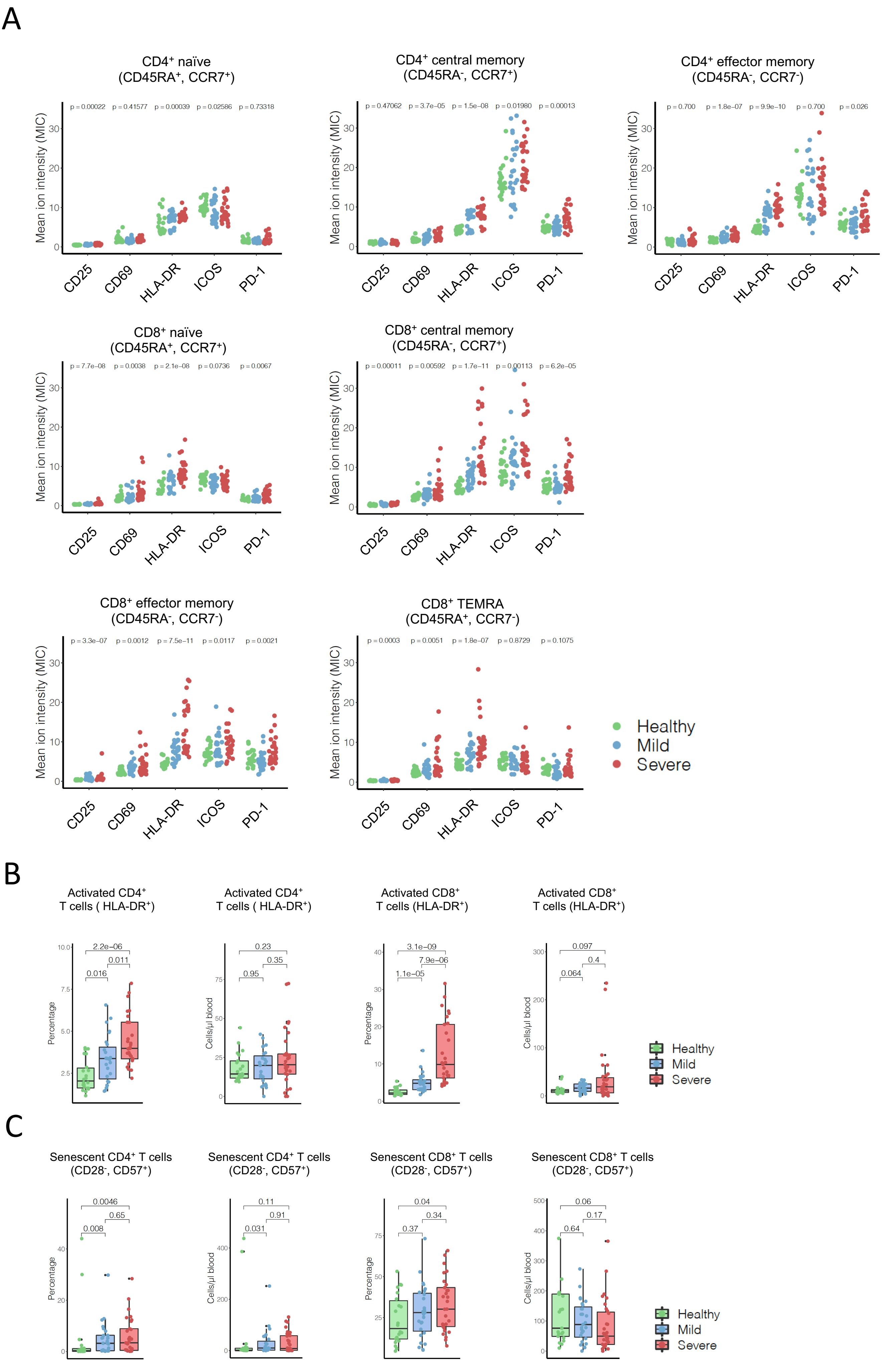


**Figure S3**. Patients with COVID-19 irrespective of disease severity have strong T cell activation.

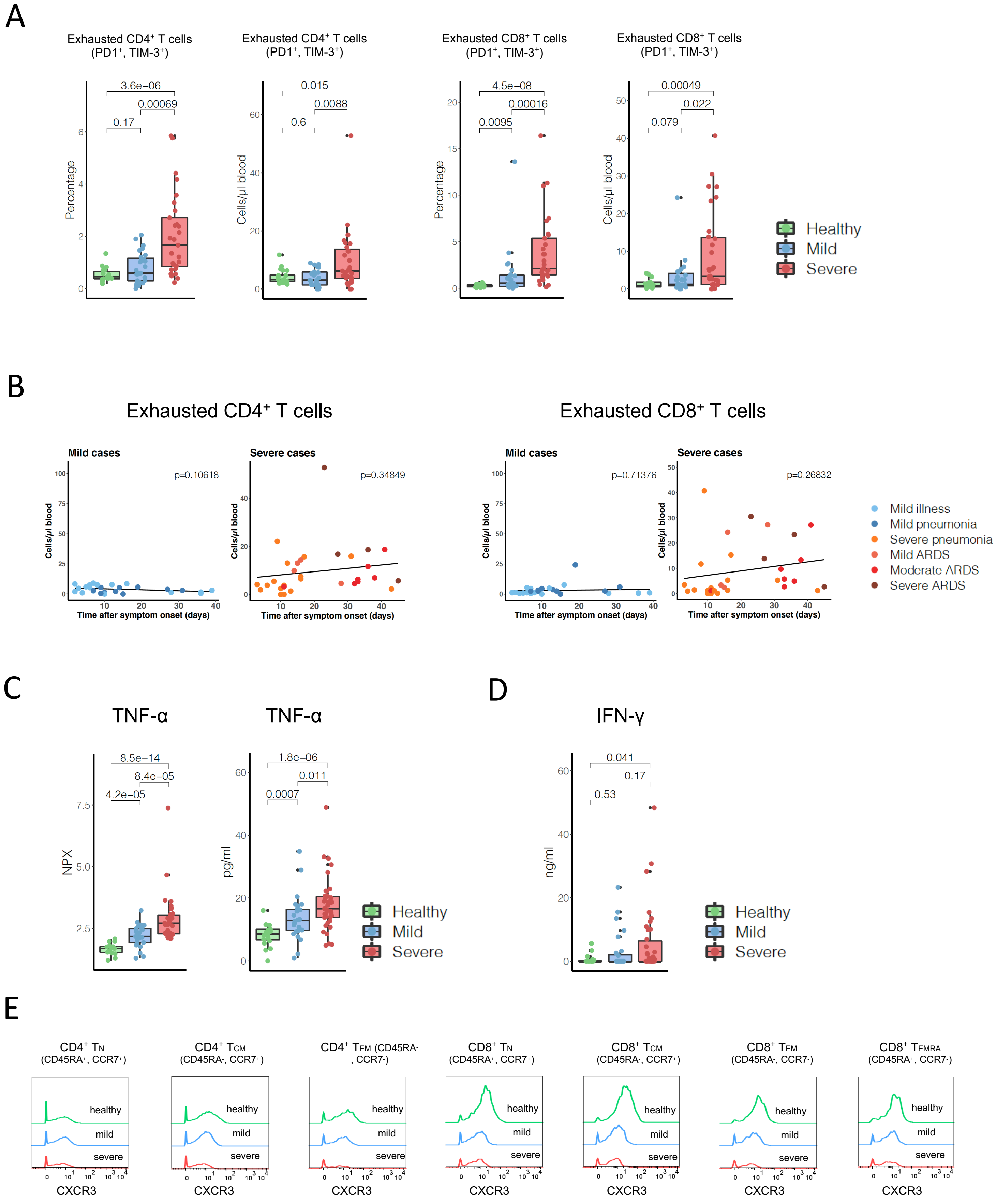


**Figure S4.** T cell exhaustion, pro-inflammatory cytokine elevation and increased T cell migration in COVID-19.

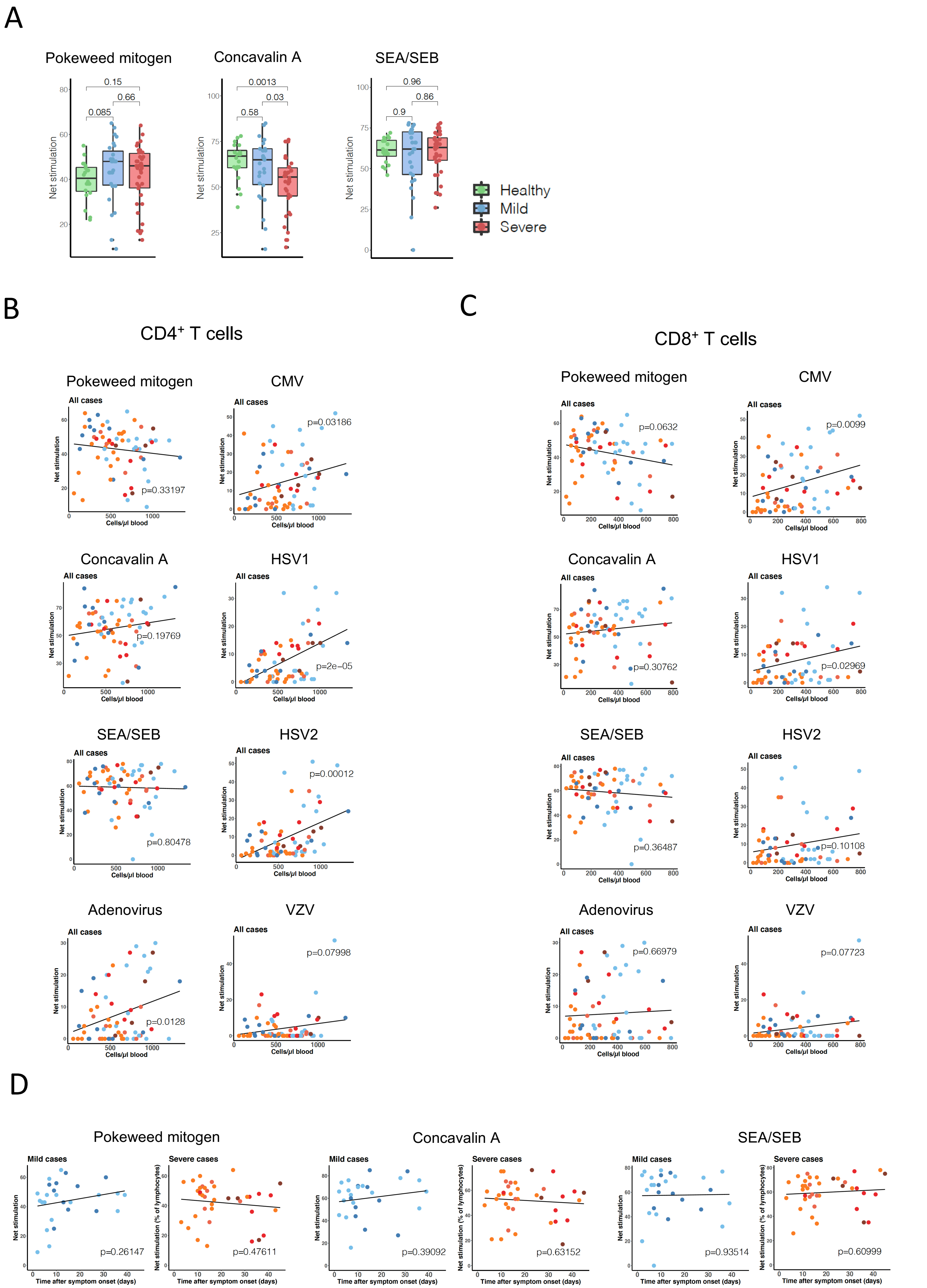


**Figure S5**. Blast formation upon stimulation with viral antigens correlates with absolute T cell counts.

A

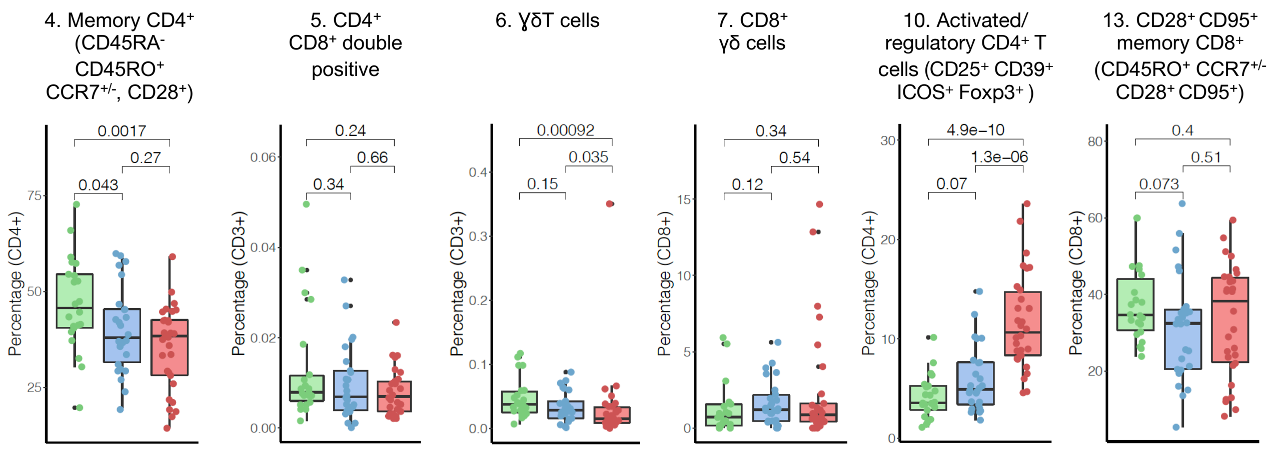


**Figure S6**. Frequencies of PhenoGraph clusters.
